# Inflammatory Profiles Induced following Intranasal Vaccination with Ricin Toxin-Immune Complexes

**DOI:** 10.1101/2023.10.03.560710

**Authors:** Lindsey E. Tolman, Nicholas J. Mantis

**Affiliations:** Department of Biomedical Sciences, School of Public Health, University at Albany, Albany, NY 12222, USA; Division of Infectious Diseases, Wadsworth Center, New York State Department of Health, Albany, NY 12208, USA

## Abstract

Ricin toxin (RT), a ribosome-inactivating protein derived from castor beans (*Ricinus communis*), is lethal in extremely small doses via inhalation, due in part to pulmonary inflammation triggered by the toxin. While no licensed countermeasures for RT exists, a myriad of toxin-neutralizing monoclonal antibodies (mAbs) against RT’s enzymatic (RTA) and binding (RTB) subunits have been described, including PB10 (anti-RTA) and SylH3 (anti-RTB). We recently demonstrated that a single administration of PB10 and SylH3 as an RT immune complex (RIC) to mice by the intranasal route functions as a potent immunogen, stimulating the rapid onset of high titer toxin-neutralizing antibodies that persist for months. In this report, we demonstrate that the de novo antibody response following RICs exposure is wholly dependent on CD4^+^ T cell help, as treatment of mice with an anti-CD4 monoclonal antibody abrogated the onset of RT-specific antibodies. To investigate other immunological factors associated with RICs, we examined the early local and systemic inflammatory cytokine and chemokine responses associated with intranasal RT and RIC administration. We report that RICs stimulate an inflammatory profile in the bronchoalveolar lavage fluids (BALF) and serum similar to that elicited by RT itself, but markedly reduced in magnitude. Of the 32 analytes examined, the most pronounced were IL-6, KC (CXCL1), G-CSF, and GM-CSF. To evaluate the role of IL-6 in driving the de novo onset of RT antibodies, groups of mice were treated (or not) with an IL-6 neutralizing monoclonal antibody prior to intranasal RICs challenge. While the monoclonal antibody treatment successfully depleted circulating IL-6 to levels below detection, it had no impact on the kinetics or magnitude of RICs-induced RT-specific antibody response. Thus, our results suggest that other cytokines/chemokines triggered by RICs are involved in influencing long lasting durable antibody responses to RT.

## INTRODUCTION

The heterodimeric glycoprotein ricin toxin (RT), synthesized as a byproduct of castor oil production in the beans of *Ricinus communis*, is an extremely potent ribosome-inactivating protein. The B subunit (RTB) binds to galactose/galactosamine residues on mammalian cells to mediate endocytic entry of the toxin, which then undergoes retrograde transport through the trans-Golgi network and endoplasmic reticulum (ER). Within the ER, the A subunit (RTA) unfolds and separates from RTB to escape into the cytoplasm, where it refolds before depurinating ribosomes, irreversibly inactivating them. This rapid ribosomal damage activates various cellular stress pathways, induces programmed cell death, and stimulates secretion of various proinflammatory cytokines. Beyond the cellular level, this combination of severe inflammation and cell death leads to extensive tissue damage and culminates in host death. Injection, ingestion, or inhalation of even small quantities of RT is generally lethal, due to both the toxin’s efficiency and the lack of any FDA-approved pre- or post-exposure therapeutics for RT. Collectively, these factors have earned RT notoriety as a Center for Disease Control and Prevention (CDC) Category B biothreat agent.

While there is currently no available preventive or resolving therapeutic options for ricin intoxication, monoclonal antibodies (mAbs) offer an excellent avenue to remedy this. The rise of mAbs as therapeutic tools for treatment of human diseases in recent years has been meteoric, with approved mAbs targeting antigens ranging from cancerous cells to live viruses. In the recent COVID-19 pandemic, mAb treatment was one of the earliest therapies available for severe cases of infection.

We have previously characterized a wide range of mAbs targeting RT, with a mix of both neutralizing and non-neutralizing activity. Among the best neutralizers are PB10, which targets RTA, and SylH3, which targets RTB. Notably, we discovered that the combination of PB10 and SylH3 together provides even greater protection against RT across *in vitro* and *in vivo* studies, as evidenced by reductions in cell/host death, inflammatory cytokine signaling, and tissue damage.

Though the protection conferred by the PB10/SylH3 mAb cocktail treatment was impressive, the efficacy of these mAbs was still believed to be limited to a relatively short term, as is generally the case for therapeutic mAb treatments. However, subsequent studies also revealed that co-administration of PB10 and SylH3 with RT to mice actually induced the rapid and prolonged generation of endogenous anti-RT IgG. This finding suggested that the use of these mAbs in combination with RT as a pre-formed immune complex held the potential to generate a humoral response akin to vaccination, opening up new avenues for both RT vaccine development as well as the expansion of therapeutic mAb applications to preventive purposes.

Additional work studying the humoral response induced by these RT-mAb immune complexes (RICs) to mice revealed that they are able to stimulate a potent humoral response against RT by both systemic (intraperitoneal; i.p.) and mucosal (intranasal; i.n.) administration routes. The ability to initiate a response by the i.n. route was made all the more impressive when we found that, compared to i.p. RICs vaccination, i.n. vaccination offered superior protection against i.n. RT challenge, and similar protection against i.p. RT challenge.

The impressive protection conferred by intranasal RICs raised interest as to the mechanisms behind RICs vaccination. While we were able to determine that RICs are able to induce potent immunity independent of FcγR, questions about the encompassing respiratory environment during the induction of immunity, including the inflammatory conditions and identification of key cytokines, remained. Further, the role of helper T (CD4^+^) cells in mediating the humoral response to RICs remained unclear. In this study we set out to define to the role of helper T cells and characterize the local and systemic inflammatory environments in the context of i.n. RICs immunization.

## MATERIALS AND METHODS

### Chemicals, biological reagents, and monoclonal antibodies (mAbs)

Ricin toxin (*Ricinus communis* agglutinin II; RCA_60_) was purchased from Vector Laboratories (Burlingame, CA, USA) and dialyzed against phosphate buffered saline (PBS) at 4°C in 10,000 MW cutoff Slide-A-Lyzer dialysis cassettes (ThermoFisher, Pittsburgh, PA, USA) prior to use in mice. Murine mAbs PB10 and SylH3 were purified by Protein G chromatography by the Dana Farber Cancer Institute’s Monoclonal Antibody Core facility [1,2]. Unless noted otherwise, all reagents were purchased from Sigma-Aldrich (St. Louis, MO, USA).

### Mouse studies

The mouse studies were conducted under strict compliance with the Wadsworth Center’s Institutional Animal Care and Use Committee (IACUC). Female BALB/c mice ages 7-8 weeks were purchased from Taconic Biosciences (Rensselaer, NY, USA). For RICs immunizations, RT (1 μg) was mixed with 20 μg each of PB10 and SylH3 and administered to mice in a final volume of 40 μl intranasally (i.n.). For RT-only exposures, 5 x LD_50_ (1 μg; unless otherwise noted) was administered to mice in a volume of 40 μl for i.n. delivery. Following RT exposure, mice were monitored daily for 7 days for symptoms of ricin intoxication, including weight loss, low blood glucose levels, and clinical signs of morbidity. Morbidity was measured using an IACUC-approved grading sheet stratified on a scale from 0-3, with 0 indicating normal activity and appearance and 3 indicating severe illness (**Table I**).

Mice were euthanized by CO_2_ asphyxiation followed by cervical dislocation when they exceeded predetermined thresholds for weight loss, blood glucose levels, or physical signs of morbidity. For studies requiring harvest of bronchoalveolar lavage fluid (BALF), mice were euthanized by CO_2_ asphyxiation and subsequent exsanguination via the abdominal aorta. For collection of BALF, 1mL of 0.6mM EDTA in PBS was used to flush the lungs intratracheally before recollection of fluid and storage at -80°C until use. Blood was procured from the submandibular vein and serum isolated as described previously [3].

### CD4^+^ T cell Depletion

Mice were intraperitoneally (i.p.) injected with 200 μg of rat anti-mouse CD4 monoclonal antibody (Clone GK1.5; ThermoFisher Scientific) 1 day prior to RICs immunization. The same dose and delivery route were used for subsequent treatment on days 1 and 4 following RICs immunization. For confirmation of antibody efficacy, mice were i.p. injected with 200 μg of rat anti-mouse CD4 monoclonal antibody 1 day before harvest of splenocytes for analysis of CD4 populations by flow cytometry in comparison to splenocytes from untreated animals.

### IL-6 Depletion

Mice were intraperitoneally (i.p.) injected with 400 μg of *InVivo*MAb anti-mouse IL-6 monoclonal antibody (Clone MP5-20F3; Bio X Cell, Lebanon, NH, USA) 1 day prior to RICs immunization. The same dose and delivery route was used for treatment 1 day after RICs immunization. For confirmation of antibody efficacy, mice were i.p. injected with 400 μg of *InVivo*MAb anti-mouse IL-6 monoclonal antibody 1 day before RICs immunization. On the day following immunization, BALF was collected for analysis of secreted IL-6 by Luminex in comparison to BALF from immunized animals that did not receive anti-mouse IL-6 treatment.

### ELISA

96-well high-binding microtiter plates (ThermoFisher Scientific) were coated overnight with ricin toxin (0.1 μg/well in PBS) at 4°C. The plates were then washed and blocked as previously described [3] before the addition of mouse sera or mAbs. Sera and mAbs were diluted in ELISA block solution (5% goat serum in 0.1% PBS-Tween). Horseradish peroxidase (HRP)-labeled, goat anti-mouse IgG-specific polyclonal antibodies (SouthernBiotech, Birmingham, AL, USA) was used as secondary reagent at a 1:2000 concentration diluted in ELISA block solution. Plates were developed using 3,30,5,50-tetramethylbenzidine (TMB; Kirkegaard & Perry Labs, Gaithersburg, MD, USA) and analyzed with a SpectraMax iD3 spectrophotometer equipped with Softmax Pro 7.1 software (Molecular Devices, Sunnyvale, CA, USA).

### Luminex

The MILLIPLEX® Mouse Cytokine/Chemokine Magnetic Bead Panel (MCYTMAG-70K-PX32) from Millipore Sigma (Burlington, MA, USA) was used to quantify cytokine contents from serum and BALF samples according to manufacturer instructions. Serum and BALF samples were diluted 1:1 into assay buffer and PBS, respectively, before combination with premixed beads on a 96-well plate for overnight incubation at 4°C with shaking. Plates were washed using a handheld plate magnet before addition of detection antibodies and 1 h of shaking incubation at RT. Streptavidin-phycoerythrin (Strep-PE) was added to each well and incubated with shaking for 30 minutes at RT. Plates were washed as described above and run in Sheath Fluid PLUS on a Luminex® FLEXMAP 3D® with INTELLIFLEX software (Luminex Corp., Austin, TX). Values were recorded as median fluorescence intensity (MFI). For analysis, assay kit standards were used to interpolate MFI values of each cytokine in each sample into a pg/mL value per manufacturer instructions before adjusting for dilution factors and normalizing to background.

### Flow cytometry

To prepare splenocyte samples, harvested spleens were collected in 10mL HBSS on ice before manual digestion through a 70 μm nylon mesh strainer. Cells were centrifuged and resuspended in lysis buffer (150mM ammonium chloride [NH_4_Cl], 0.7mM potassium phosphate monobasic [KH_2_PO_4_] in dH_2_O) for 4 minutes at RT with agitation. HBSS was added to samples before centrifugation, counting, and resuspension at 5x10^6^ cells/mL in ice cold FACS buffer (1% FBS, 1mM EDTA in PBS).

For all experiments, 500 uL cell suspension was added to 12x75mm polystyrene tubes and incubated with 100 uL TruStain FcX™ (BioLegend, San Diego, CA, USA) on ice for 20 min. The following antibodies were used for labeling cells: PE rat anti-mouse CD45 (Clone 30-F11; BioLegend), FITC rat anti-mouse CD3 (Clone 17A2; BD Biosciences, San Jose, CA, USA), and APC rat anti-mouse CD4 (Clone RM4-5; BD Biosciences). 50 uL of each antibody was added to samples and incubated on ice in the dark for 1 h. Cells were fixed using BD Cytofix™ Fixation Buffer (BD Biosciences) according to manufacturer instructions and stored at 4°C protected from light before acquisition on a FACSCalibur cytometer equipped with CellQuest Pro software (BD Biosciences) within 1 week of fixation. Analysis was performed using FlowJo v10.8.0 (BD Biosciences).

### Statistical analyses

Statistical analyses of weights and titers were carried out using GraphPad Prism 9.2.0. Unpaired, two-tailed Welch’s t-tests were performed to determine the significance of differences in end-point titers between groups as well as in IL-6 contents in BALF and serum. In all cases, * p ≤ 0.05, ** p ≤ 0.01, *** p ≤ 0.001, **** p < 0.0001.

## RESULTS

### Role of CD4^+^ T cells in stimulating de novo RT-specific IgG following intranasal RICs immunization

We recently reported that intranasal administration of RICs to mice induces a rapid and durable serum anti-RT IgG response within 10-14 days without a detectable IgM phase [3]. To ascertain whether CD4^+^ T cells are involved in this response, groups of adult BALB/c mice were treated with anti-CD4 antibodies (400 μg by intraperitoneal injection) before and after intranasal RICs administration on study day 0. The anti-CD4 antibody regimen resulted in a near complete of depletion of CD4^+^ T cells, as determined in a parallel group of animals (**Figure 1A, B**). RT-specific serum IgG titers were measured on days 7, 14 and 30 following RICs administration.

**Figure 1.**
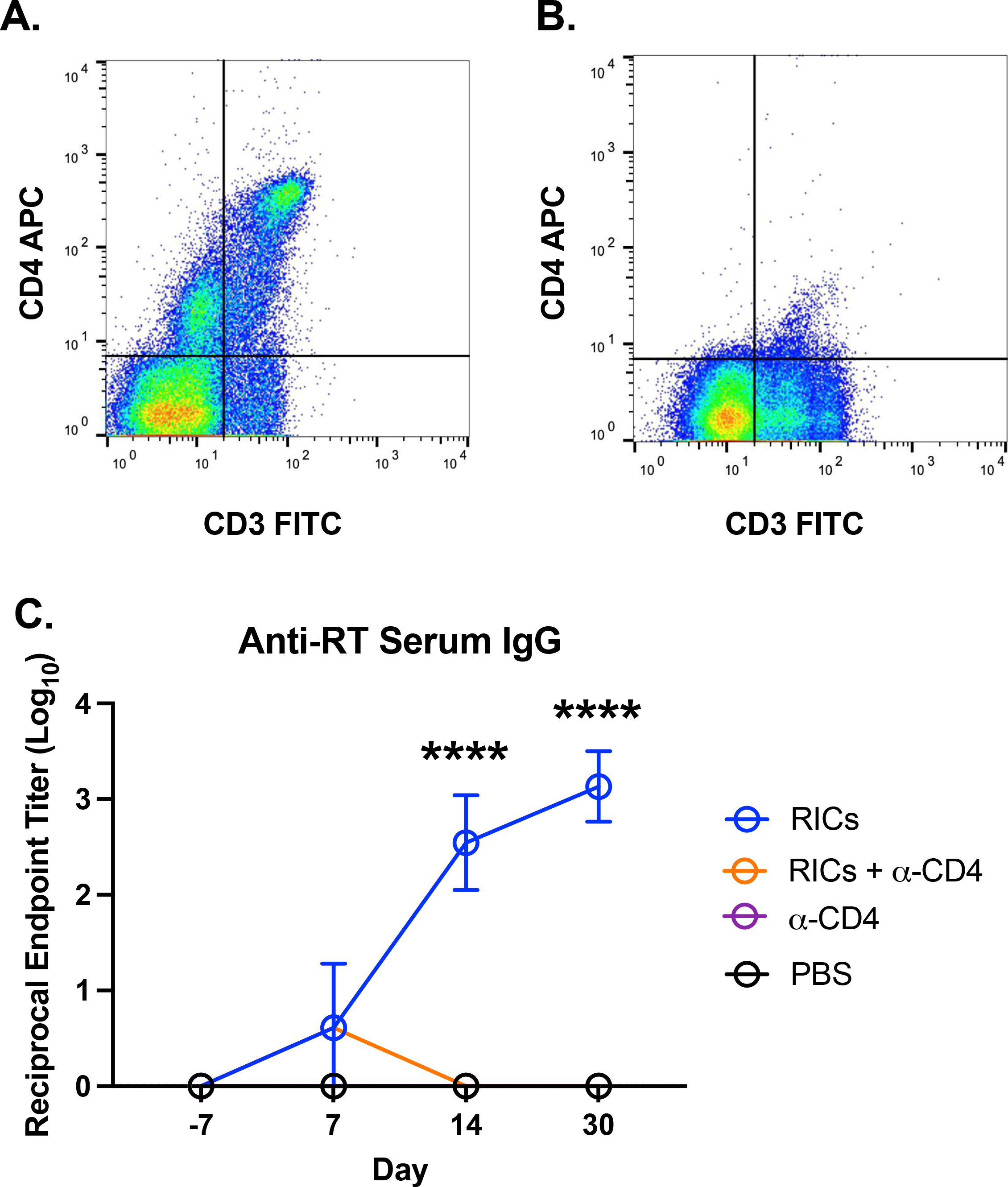
Helper T cells are required for mounting a humoral response following i.n. RICs immunization. RICs were administered i.n. alone or in tandem with anti-CD4 treatment. (**A**) Combined population of helper T cells (CD3^+^CD4^+^) in the spleens of untreated mice (n=3). (**B**) Combined population of helper T cells in the spleens of mice treated with anti-CD4 antibody (n=3). (**C**) Anti-RT serum IgG titers of RICs-immunized mice receiving no additional treatment or anti-CD4 antibody (n=6), or control animals receiving only anti-CD4 antibody or PBS (n=4). Significance of endpoint titers was determined by unpaired, two-tailed Welch’s t-test.

In the control group of animals, intranasal RICs administration resulted in the onset of RT-specific serum IgG by day 14, with reciprocal endpoint titers exceeding 10^3^ by day 30 (**Figure 1C**). In contrast, mice treated with anti-CD4 antibodies had RT-specific IgG levels that were essentially at baseline, even on day 30 (**Figure 1C**). This result demonstrates the necessity of CD4^+^ T cells in the humoral response to RICs.

### Induction of mucosal and systemic inflammatory cytokines and chemokines following RICs exposure

We next sought to investigate what impact RIC exposure has on local inflammatory cytokine and chemokines levels, reasoning that the nature of the response may provide a clue as to the underlying factors that drive durable anti-toxin B cell responses [4,5]. To address this question, we collected serum and bronchoalveolar lavage fluids (BALFs) from mice 6, 12, and 18 h after intranasal RIC (20μg PB10, 20μg SylH3, 1μg RT) administration, and then subjected samples to a 32-plex mouse cytokine/chemokine array. For comparison, we also collected samples from groups of mice treated i.n. with RT (1μg), the PB10/SylH3 mAb cocktail (20μg each mAb) or PBS. Neither PBS nor mAb treatments induced a notable change in inflammatory cytokine/chemokines in either serum or BALF (**Supplemental Table I**).

Intranasal delivery of RT stimulated the secretion of a variety of proinflammatory cytokines in serum and BALF. By 6 h after administration, several cytokines were moderately increased in BALF compared to control animals, while 14 of the 32 tested cytokines in the serum were slightly elevated (**Figure 2**). This early uptick in cytokine production is indicative of the burgeoning inflammatory response that quickly follows RT exposure. This trend continues at the 12 h time point, when inflammatory cytokines in the BALF reach peak concentrations. Between the 6 and 12 h time points, average BALF concentrations of G-CSF, LIF, KC (CXCL1), MIP-1β, and MIP-2 grew roughly 5-fold, IL-1β, IL-6, MIP-1α grew approximately 10-fold, and GM-CSF and VEGF grew about 20-fold in respect to the mean concentrations of each measured at the 6 h mark (**Table I**). Cytokine levels remain high in BALF at the 18 h mark as well. This enormous production increase in about 1/3 of tested cytokines only 12 h after pulmonary RT exposure, and their subsequent maintenance at high concentrations, demonstrates just how inflammatory RT is in the respiratory tract.

**Figure 2.**
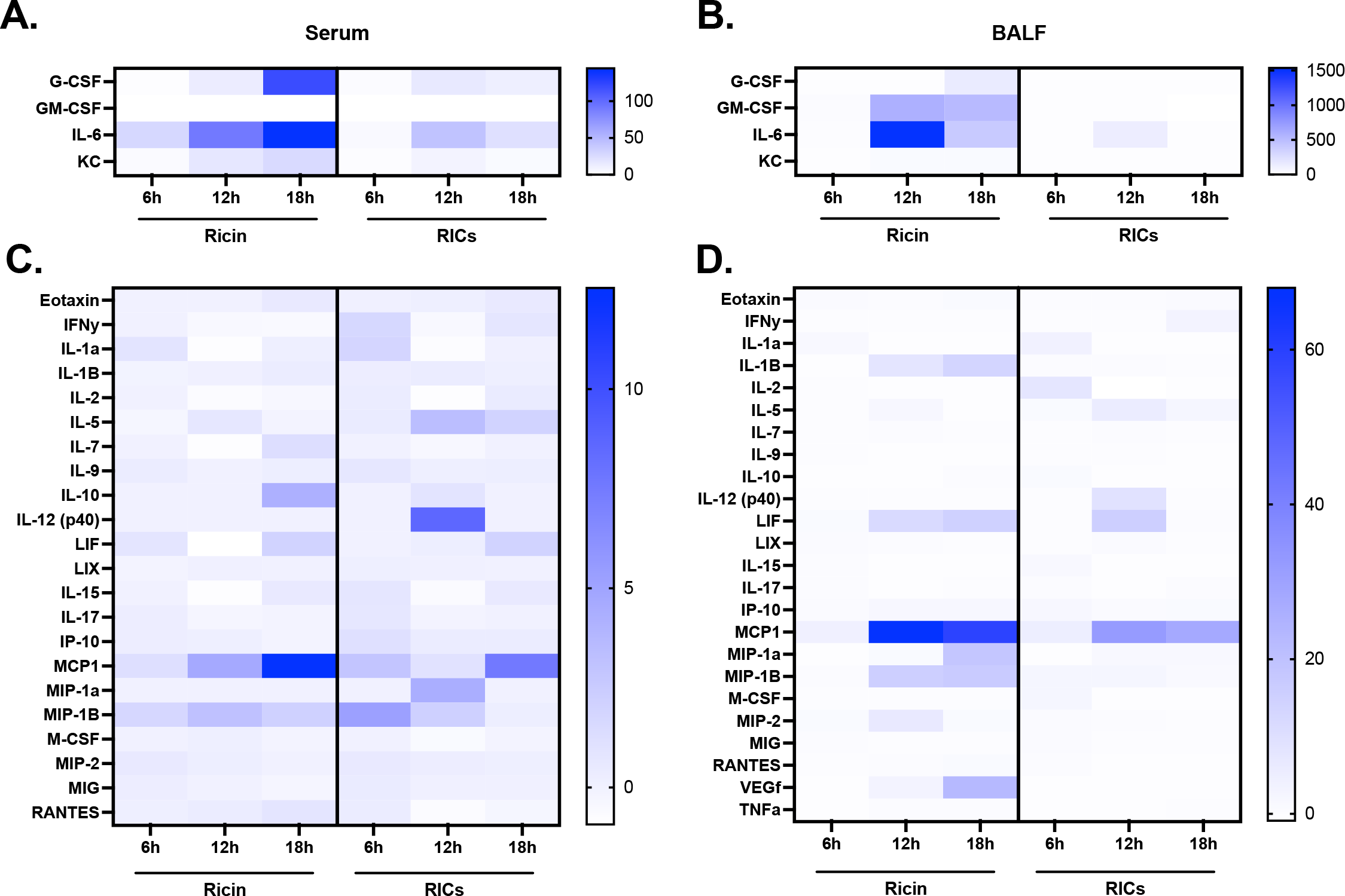
RICs induce an intermediate inflammatory environment within the respiratory tract. RICs, RT, or PBS were administered to mice by the i.n. route before harvest of serum and BALF at 6, 12, or 18 h post-treatment for cytokine content analysis. For each treatment at each time point, n=6. Concentrations are expressed as fold-change over PBS-treated samples. Notable inflammatory cytokines produced in high quantities in both (**A**) serum and (**B**) BALF. Cytokines with a nonzero fold change at any tested time point are shown in (**C**) for serum samples and (**D**) for BALF samples. Scales for fold change heatmaps in panels **A-D** vary. Mean group concentration values for all samples, including those with zero fold change, and their statistical significance relative to control samples can be found in **Supplemental Table I**. Radical variation was seen in the concentration of some cytokines, compared to others in the panel, based on treatment and sample source. Special attention should be given to the varying scales associated with the heatmaps of Figure 2 that reflect these differences.

RT induces a strong inflammatory response in serum at 12 and 18 h post-exposure, though their kinetics and magnitude vary from the trends seen in BALF. While cytokine levels are elevated in serum compared to control animals, they only reach roughly 20% or less of the levels of their respective counterparts in BALF throughout the 18 h observed in this study (**Table I**). Interestingly, though BALF cytokine concentrations peaked at the 12 h mark, in serum they continued to gradually rise across all subsequent time points, with their greatest magnitudes at 18 h post-exposure (**Figure 2A, C**).

Particularly elevated cytokines following RT exposure included KC (CXCL1) in serum, GM-CSF in BALF, and IL-6 and G-CSF in both serum and BALF (**Figure 2A, B**). IL-6, possibly the most signature cytokine associated with RT inflammation, was by far the most pronounced in terms of fold change. This profile agrees with other reports that have examined the impact of RT on lung inflammation in mice and NHPs [6–9].

RICs also stimulated an inflammatory response in the BALF, with important deviations from RT-treated animal samples. Interestingly, the cytokines produced in greater quantity following RT exposure were generally the same ones elevated by RICs delivery, though RICs induced a markedly less inflammatory response than RT at all time points. A clear example of this trend is the quantity of IL-6 in BALF 12 h post-treatment. In RT samples at this time point, average IL-6 concentration was 1717.61 pg/mL, over 1500-fold greater than the concentration found in matched PBS samples. In contrast, IL-6 in RICs samples at this time point averaged a concentration of 153.04 pg/mL, which was elevated roughly 135-fold over PBS sample levels but still under 1000-fold less than the level induced by RT. Also similar to RT, inflammatory cytokines in RICs BALF samples were modestly increased at the 6 h mark, peaked at 12 h post-administration, and slightly dwindled by the 18 h point.

In serum samples, RICs again induced inflammation via similar cytokines and at a lower intensity than that seen in RT samples. As observed in RT samples, RICs also induced much lower inflammation in the serum than in the BALF. However, RICs stimulated a more complex pattern of cytokine concentration in the serum. While RT samples gradually rose in concentration, peaking at the 18 h mark, RICs samples did not portray an overall increasing or decreasing systemic inflammation trend. Some cytokines, such as MIP-1β, peaked at 6 h post-administration, while others like IL-12 (p40) and MCP1 peaked at 12 and 18 h, respectively.

This paints a picture of a much more complicated management of inflammation in the serum, suggesting that RICs induce a well-regulated systemic response that remains balanced over time and refrains from reaching extreme levels of inflammation.

This cytokine profiling of i.n. RICs immunization highlights the increased production of G-CSF, KC, MIP-1α, and MIP-1β in serum (**Figure 2A, C**), as well as MCP1 and IL-6 in both BALF and serum. The increased production of IL-6 following pulmonary delivery of both RT and RICs is particularly interesting, as IL-6 is a key mediator of multiple diseases, including rheumatoid arthritis (RA), Castleman disease, COVID-19, and cytokine release syndrome (CRS) [10,11], the formal name associated with cytokine storms such as those seen following RT exposure.

These findings demonstrate that the early inflammation that follows i.n. RICs immunization is greatly dampened relative to that induced by pulmonary RT exposure. This data also suggests that the early inflammatory response following i.n. RICs immunization is mediated primarily within the lungs, based on changes in cytokine concentration, thus creating a controlled local inflammatory environment suitable for initiation of a humoral response.

### Local IL-6 signaling is not critical for mediation of a humoral response to RICs

IL-6 was notably pronounced in BALF and serum of mice following intranasal RICs exposure. As IL-6 has numerous (direct and indirect) roles in promoting B cell responses, we postulated that it might contribute to the observed onset of RT-specific IgG titers following RICs treatment. Our determination of the necessity of helper T cells in mediating the humoral response to RICs also drew attention to the heightened IL-6 levels, as other work has highlighted the relationship between IL-6 signaling and T cell responses [12,13].

To investigate this possibility, we first examined whether treatment of BALB/c mice with an IL-6 neutralizing mAb was sufficient to deplete IL-6 levels following RICs treatment. Indeed, IL-6 levels one day following RICs exposure were below the level of detection in serum and significantly reduced (but not fully eliminated) in BALF, as compared to control mice (**Figure 3A, B**). Next, groups of mice were treated (or not) with the IL-6 neutralizing mAb 1 day before and 1 day after intranasal RICs exposure and then monitored for the onset of RT-specific IgG over the course of 30 days. The kinetics and magnitude of RT-specific IgG titers following RICs exposure were identical in mice that received IL-6 neutralizing antibody, as compared to the control animals (**Figure 3C**). These findings suggest that IL-6 is dispensable for modulating the humoral response to RICs, although subject to the caveat that IL-6 neutralization may not have fully penetrated the lung and we cannot completely exclude the possibility that even locally reduced levels of IL-6 are sufficient to promote antibody onset.

**Figure 3.**
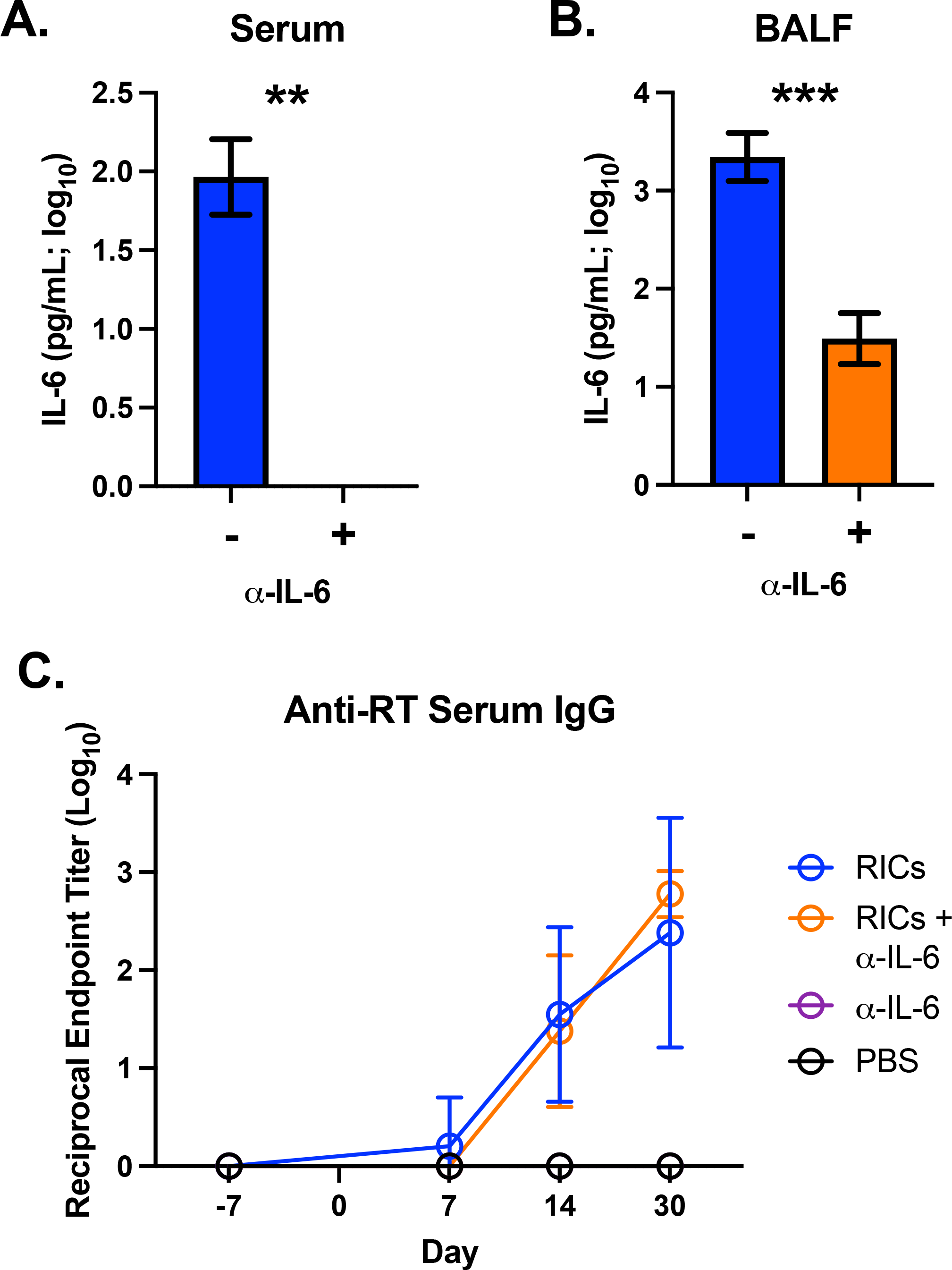
IL-6 signaling is not required for i.n. RICs vaccination. Adult mice were treated i.p. with anti-mouse IL-6 mAb 1 day prior to i.n. RICs immunization. (**A**) Serum and (**B**) BALF levels of IL-6 from naїve (n=3) and (**C, D**) anti-mouse IL-6-treated animals (n=3) 18 h after i.n. RICs administration. (**E**) Anti-RT Serum IgG titers of treated and untreated animals following i.n. RICs immunization. Significance of endpoint titers was determined by unpaired, two-tailed Welch’s t-test.

## DISCUSSION

The encompassing environment is highly influential in determining if and how an adaptive immune response is mounted against an antigen. While some tissues, such as those of the lymphoid system, are poised to mediate adaptive responses, mucosal tissues such as the respiratory tract generally resist reactivity and tend toward antigenic tolerance. Antigens that are able to overcome this mucosal tolerance, such as RT, often cause extensive inflammation that is ultimately more damaging than beneficial for the generation of long-term protective immunity. In contrast, RICs delivered i.n. to mice stimulate a rapid and considerable humoral response with little to no negative clinical impact. Here we sought to characterize the inflammatory setting associated with i.n. RICs immunization to better understand the early local and systemic environments in which RICs successfully stimulate protective immunity within the respiratory tract.

While our recent work with RICs demonstrated the induction of a strong humoral response, the role of T cells in mediating this immunity remained unclear. Here we demonstrate by depletion of CD4^+^ cells that helper T cells are indeed involved in the induction of humoral immunity to i.n. RICs vaccination. Further, the loss of helper T cells stunted typical IgG production, indicating that CD4^+^ cells are not only involved in the process of i.n. RICs vaccination, but are essential to the process. We then explored the broader inflammatory response following RICs immunization, finding that they induce a similar inflammatory pattern to RT, though to a much more attenuated degree. The most dramatic differences in inflammation were seen in the BALF, in which RT exposure resulted in extremely high proinflammatory cytokine production above baseline values, while that induced by RICs was markedly dampened by comparison. In both serum and BALF samples, the greatest differences observed between the two treatments were seen in concentrations of IL-6. However, neutralization of IL-6 did not dampen the humoral response, indicating that it is not functionally essential for mounting adaptive immunity to RICs immunization.

Prior to this work, the role of T cell help in mediating the humoral response to RICs was unclear. While the secretion of anti-RT IgG indicated B cell class-switching and hinted to T cell involvement, the rapid antibody response was more characteristic of a T-independent response. We now show that CD4^+^ cells are not only involved, but essential for mounting a humoral response to RICs vaccination. The necessity of T cell help is particularly interesting, as previous work reported endogenous IgG generation within roughly 7 days of i.n. RICs administration [3]. This rapid timeline for a T-dependent adaptive response is unusual, as T cell activation alone can take 24 h [14], and germinal centers (GCs) typically do not begin forming until approximately 4 days after initial antigen exposure [15], with full formation about 7-10 days after antigen encounter [16]. Thus, i.n. administration of RICs stimulates an extremely fast T-dependent humoral response.

This abnormity calls into question where RICs antigen is actually being presented. Canonically, APCs migrate to the draining lymph node (dLN) to present processed antigen to T cells. However, a growing body of literature has demonstrated in situ antigen presentation, wherein antigen presentation and adaptive response development occur within the lungs themselves [17–19]. Under these conditions, the timeline for an adaptive response would be shortened, which would explain how RICs induce such a quick humoral response in the respiratory tract. Though the involvement of local antigen presentation of RICs is plausible, it is currently unclear and we are continuing work investigating its role in i.n. RICs vaccination.

The results presented here comprise the first inflammatory profiling of RICs, though they are not the first to characterize the cytokine response to pulmonary RT exposure. Multiple studies of aerosolized RT exposure in NHP models have been published [6,9], and the most upregulated cytokines in BALF and serum (IL-6, MCP1, MIP-1α, MIP-1β) reported in these studies are reflected by our findings in the murine model. To our knowledge, only two prior studies have presented cytokine data following pulmonary RT exposure in mice. Both were limited in scope, measuring concentrations of 5 or fewer cytokines [7,8]. Moreover, all of the discussed studies only monitored cytokine concentrations at a single time point, with 24 h post-exposure being the earliest reported. The data presented in this paper provides expanded insight into the inflammatory nature of both RT and RICs at early time points after administration, and presents findings on the largest pulmonary RT inflammation panel to date.

Collectively, these results suggest that i.n. RICs vaccination requires controlled inflammation, especially within the respiratory tract, as opposed to the excessive inflammation caused by delivery of RT alone. While the cytokine production following RICs administration largely mirrored that seen in RT-treated samples, the quantity of cytokines in RICs samples was notably lower, particularly in the case of strong chemoattractants such as KC and proinflammatory signalers like IL-6. These results demonstrate that a moderately increased inflammatory profile primarily in the respiratory tract is a key feature of i.n. RICs immunization. However, depletion of IL-6 did not impair the adaptive response to RICs, suggesting that the inflammation that follows RICs delivery may primarily serve to recruit immune cells and set the stage for an adaptive response.

Defining the environmental conditions that allow for successful intranasal immunization is an important step forward in the development of mucosal vaccines. The work presented here highlights the role of a controlled inflammatory setting in locales where adaptive responses are being developed, particularly in the context of the respiratory tract. As an increasing number of studies demonstrate the benefits of mucosal vaccination for improved protection against a myriad of agents, it is essential that we continue to elucidate the prime conditions for inducing long-term local immunity.

## ACKNOWLEDGMENTS

We gratefully acknowledge members of the Wadsworth Center’s Veterinary Sciences team for caring for our animals. We would also like to thank Dr. Renjie Song of the Wadsworth Center’s Immunology Core for assistance in flow cytometry experiments and Grace Freeman-Gallant for her guidance in the completion of the Luminex assays presented here.

## FOOTNOTES

1 This study was supported by grant AI125190 and Contract No. HHSN272201400021C from the National Institute of Allergy and Infectious Diseases (NIAID), National Institutes of Health (NIH). The content is solely the responsibility of the authors and does not necessarily represent the official views of the NIH. The funders had no role in study design, data collection and analysis, decision to publish, or preparation of the manuscript.

