## Supplemental Table 1 for "Inflammatory Profiles Induced following Intranasal Vaccination with Ricin Toxin-Immune Complexes"

| Table I. Cytokine Concentrations and Fold Changes | | | | | | | | |
|---|---|---|---|---|---|---|---|---|
| Cytokine | Treatment | Source | pg/mL[a,b] | | | Fold Change[b,c] | | |
| | | | 6h | 12h | 18h | 6h | 12h | 18h |
| G-CSF | RT | Serum | 351.87 | 1738.71* | 11032.59** | 0.57 | 13.38 | 119.03 |
| | | BALF | 380.10 | 2251.47* | 12954.94** | 0.74 | 17.83 | 155.14 |
| | RICs | Serum | 840.17 | 2053.47 | 1154.41 | 2.75 | 15.98 | 11.56 |
| | | BALF | 833.95 | 2119.11 | 1214.11 | 2.81 | 16.72 | 13.63 |
| | mAbs | Serum | 1272.48 | 1227.35 | 449.77 | 4.69 | 9.15 | 3.89 |
| | | BALF | 596.48 | 613.02 | 181.14 | 1.72 | 4.13 | 1.18 |
| | PBS | Serum | 223.81 | 120.94 | 91.91 | 0 | 0 | 0 |
| | | BALF | 218.93 | 119.56 | 82.97 | 0 | 0 | 0 |
| Eotaxin | RT | Serum | 367.71 | 312.31 | 441.24 | -0.03 | -0.02 | 0.54 |
| | | BALF | 310.62 | 226.71 | 313.49 | 0.68 | 0.30 | 0.79 |
| | RICs | Serum | 390.34 | 365.19 | 421.74 | 0.03 | 0.14 | 0.47 |
| | | BALF | 250.72 | 219.15 | 261.64 | 0.36 | 0.26 | 0.50 |
| | mAbs | Serum | 398.21 | 330.09 | 318.15 | 0.05 | 0.03 | 0.11 |
| | | BALF | 217.72 | 121.66 | 55.11 | 0.18 | -0.30 | -0.68 |
| | PBS | Serum | 377.55 | 319.15 | 286.62 | 0 | 0 | 0 |
| | | BALF | 184.83 | 172.80 | 174.71 | 0 | 0 | 0 |
| GM-CSF | RT | Serum | 0 | 0 | 0 | 0 | 0 | 0 |
| | | BALF | 25.87 | 587.20 | 518.85 | 24.87 | 586.20 | 517.85 |
| | RICs | Serum | 0 | 0 | 0 | 0 | 0 | 0 |
| | | BALF | 14.37 | 9.56 | 0 | 13.37 | 8.56 | 0 |
| | mAbs | Serum | 0 | 0 | 0 | 0 | 0 | 0 |
| | | BALF | 14.26 | 9.96 | 1.91 | 13.26 | 8.96 | 0.91 |
| | PBS | Serum | 0 | 0 | 0 | 0 | 0 | 0 |
| | | BALF | 0 | 0 | 0 | 0 | 0 | 0 |
| IFNγ | RT | Serum | 0 | 0 | 0 | 0 | -0.48 | -0.56 |
| | | BALf | 0 | 0 | 0 | 0 | 0 | 0 |
| | RICs | Serum | 2.71 | 0 | 3.93 | 1.71 | -0.48 | 0.74 |
| | | BALF | 0 | 0 | 3.93 | 0 | 0 | 2.93 |
| | mAbs | Serum | 0 | 0 | 0 | 0 | -0.48 | -0.56 |
| | | BALF | 0.92 | 1.18 | 1.23 | -0.08 | 0.18 | 0.23 |
| | PBS | Serum | 0 | 1.93 | 2.27 | 0 | 0 | 0 |
| | | BALF | 1.00 | 1.00 | 1.00 | 0 | 0 | 0 |
| IL-1α | RT | Serum | 177.89 | 134.62 | 142.30 | 0.91 | -0.78 | 0.18 |

| | | | | | | | | |
|---|---|---|---|---|---|---|---|---|
| | | BALF | 152.85 | 108.71 | 105.99 | 1.44 | -0.82 | -0.09 |
| | RICs | Serum | 266.73 | 160.01 | 126.61 | 1.86 | -0.74 | 0.05 |
| | | BALF | 283.35 | 117.36 | 81.24 | 3.53 | -0.81 | -0.31 |
| | mAbs | Serum | 170.01 | 226.70 | 150.13 | 0.82 | -0.63 | 0.24 |
| | | BALF | 85.08 | 50.64 | 26.06 | 0.36 | -0.92 | -0.78 |
| | PBS | Serum | 93.24 | 611.23 | 120.78 | 0 | 0 | 0 |
| | | BALF | 62.56 | 609.32 | 117.10 | 0 | 0 | 0 |
| IL-1β | RT | Serum | 0.90 | 1.09 | 1.22 | -0.13 | 0.06 | 0.36 |
| | | BALF | 0.90 | 9.06 | 13.05 | -0.10 | 8.06 | 13.54 |
| | RICs | Serum | 1.32 | 1.42 | 1.03 | 0.28 | 0.38 | 0.15 |
| | | BALF | 1.32 | 1.42 | 1.03 | 0.32 | 0.42 | 0.15 |
| | mAbs | Serum | 1.29 | 1.03 | 1.00 | 0.26 | 0 | 0.11 |
| | | BALF | 7.99 | 7.81 | 6.15 | 6.99 | 6.81 | 5.85 |
| | PBS | Serum | 1.03 | 1.03 | 0.90 | 0 | 0 | 0 |
| | | BALF | 1.00 | 1.00 | 0.90 | 0 | 0 | 0 |
| IL-2 | RT | Serum | 0 | 1.64 | 1.39 | -0.04 | -0.68 | -0.43 |
| | | BALf | 0 | 1.64 | 1.39 | -0.04 | -0.86 | -0.78 |
| | RICs | Serum | 1.43 | 1.13 | 3.46 | 0.37 | -0.78 | 0.42 |
| | | BALF | 9.07 | 1.13 | 3.46 | 7.72 | -0.90 | -0.46 |
| | mAbs | Serum | 4.42 | 7.78 | 3.79 | 3.25 | 0.54 | 0.55 |
| | | BALF | 7.99 | 7.81 | 6.15 | 6.99 | 6.81 | 5.85 |
| | PBS | Serum | 1.04 | 5.05 | 2.44 | 0 | 0 | 0 |
| | | BALF | 1.00 | 1.00 | 0.90 | 0 | 0 | 0 |
| IL-4 | RT | Serum | 0 | 0 | 0 | 0 | 0 | 0 |
| | | BALF | 0 | 0 | 0 | 0 | 0 | 0 |
| | RICs | Serum | 0 | 0 | 0 | 0 | 0 | 0 |
| | | BALF | 0 | 0 | 0 | 0 | 0 | 0 |
| | mAbs | Serum | 0 | 0 | 0 | 0 | 0 | 0 |
| | | BALF | 5.57 | 3.54 | 1.26 | 4.35 | -0.69 | -0.80 |
| | PBS | Serum | 0 | 0 | 0 | 0 | 0 | 0 |
| | | BALF | 1.04 | 11.31 | 6.39 | 0 | 0 | 0 |
| IL-3 | RT | Serum | 0 | 0 | 0 | 0 | 0 | 0 |
| | | BALF | 0 | 0 | 0 | 0 | 0 | 0 |
| | RICs | Serum | 0 | 0 | 0 | 0 | 0 | 0 |
| | | BALF | 0 | 0 | 0 | 0 | 0 | 0 |
| | mAbs | Serum | 0 | 0 | 0 | 0 | 0 | 0 |

| | | | | | | | | |
|---|---|---|---|---|---|---|---|---|
| | | BALF | 0.85 | 0.87 | 0.70 | -0.15 | -0.13 | -0.30 |
| | PBS | Serum | 0 | 0 | 0 | 0 | 0 | 0 |
| | | BALF | 1.00 | 1.00 | 1.00 | 0 | 0 | 0 |
| IL-5 | RT | Serum | 22.39 | 15.28 | 4.31 | -0.33 | 0.70 | -0.21 |
| | | BALF | 16.59 | 16.79 | 5.82 | -0.03 | 1.67 | -0.11 |
| | RICs | Serum | 48.34 | 39.36 | 16.40 | 0.44 | 3.39 | 1.99 |
| | | BALF | 33.52 | 42.45 | 19.05 | 0.96 | 5.75 | 1.92 |
| | mAbs | Serum | 50.44 | 22.75 | 9.27 | 0.50 | 1.54 | 0.69 |
| | | BALF | 9.93 | 6.85 | 3.62 | -0.42 | 0.09 | -0.45 |
| | PBS | Serum | 33.60 | 8.97 | 5.49 | 0 | 0 | 0 |
| | | BALF | 17.08 | 6.29 | 6.55 | 0 | 0 | 0 |
| IL-6 | RT | Serum | 43.37 | 91.26*** | 145.53**** | 27.87 | 90.26 | 144.53 |
| | | BALF | 196.28** | 1717.61** | 2349.98*** | 20.50 | 1536.38 | 400.67 |
| | RICs | Serum | 8.15 | 43.56 | 22.99 | 4.43 | 42.56 | 21.99 |
| | | BALF | 54.04 | 153.04* | 80.56** | 4.92 | 135.98 | 12.77 |
| | mAbs | Serum | 31.56 | 10.76 | 10.73 | 20.01 | 9.76 | 9.73 |
| | | BALF | 466.44 | 309.53 | 50.78 | 50.09 | 276.05 | 7.68 |
| | PBS | Serum | 1.50 | 1.00 | 1.00 | 0 | 0 | 0 |
| | | BALF | 9.13 | 1.12 | 5.85 | 0 | 0 | 0 |
| IL-7 | RT | Serum | 0 | 0 | 2.25 | 0 | -0.88 | 1.25 |
| | | BALF | 0 | 0 | 0 | 0 | 0.20 | 0 |
| | RICs | Serum | 0 | 4.95 | 0 | 0 | -0.40 | 0 |
| | | BALF | 0 | 0 | 0 | 0 | 0.20 | 0 |
| | mAbs | Serum | 1.00 | 1.00 | 0.84 | 0 | -0.88 | -0.16 |
| | | BALF | 1.00 | 1.00 | 1.00 | 0 | 0.20 | 0 |
| | PBS | Serum | 1.00 | 8.29 | 1.00 | 0 | 0 | 0 |
| | | BALF | 1.00 | 0.84 | 1.00 | 0 | 0 | 0 |
| IL-9 | RT | Serum | 88.91 | 79.88 | 73.64 | 0.36 | 0.01 | 0.19 |
| | | BALF | 149.27 | 125.90 | 161.97 | 0.16 | -0.33 | -0.21 |
| | RICs | Serum | 105.65 | 87.66 | 74.07 | 0.62 | 0.11 | 0.20 |
| | | BALF | 151.09 | 185.96 | 121.28 | 0.17 | -0.01 | -0.41 |
| | mAbs | Serum | 107.41 | 75.56 | 72.80 | 0.65 | -0.04 | 0.18 |
| | | BALF | 149.19 | 164.07 | 153.60 | 0.16 | -0.12 | -0.25 |
| | PBS | Serum | 65.21 | 78.84 | 61.86 | 0 | 0 | 0 |
| | | BALF | 129.12 | 187.28 | 205.39 | 0 | 0 | 0 |
| IL-10 | RT | Serum | 0 | 0 | 5.23 | 0.01 | 0 | 4.23 |

| | | | | | | | | |
|---|---|---|---|---|---|---|---|---|
| | | BALF | 0 | 2.02 | 7.13 | -0.17 | -0.80 | 0.38 |
| | RICs | Serum | 0 | 1.88 | 0 | 0.01 | 0.88 | 0 |
| | | BALF | 2.26 | 1.88 | 0 | 0.88 | -0.81 | -0.81 |
| | mAbs | Serum | 1.00 | 1.00 | 1.00 | 0.01 | 0 | 0 |
| | | BALF | 1.53 | 1.18 | 1.20 | 0.27 | -0.88 | -0.77 |
| | PBS | Serum | 0.99 | 1.00 | 1.00 | 0 | 0 | 0 |
| | | BALF | 1.20 | 9.90 | 5.18 | 0 | 0 | 0 |
| IL-12 (p40) | RT | Serum | 0 | 0 | 0 | 0 | 0 | 0 |
| | | BALF | 0 | 0 | 0 | 0 | 0 | 0 |
| | RICs | Serum | 0 | 9.62 | 0 | 0 | 8.62 | 0 |
| | | BALF | 0 | 9.62 | 0 | 0 | 8.62 | 0 |
| | mAbs | Serum | 1.00 | 1.00 | 1.00 | 0 | 0 | 0 |
| | | BALF | 1.00 | 1.00 | 1.95 | 0 | 0 | 0.95 |
| | PBS | Serum | 1.00 | 1.00 | 1.00 | 0 | 0 | 0 |
| | | BALF | 1.00 | 1.00 | 1.00 | 0 | 0 | 0 |
| IL-12 (p70) | RT | Serum | 0 | 0 | 0 | 0 | 0 | 0 |
| | | BALF | 0 | 0 | 0 | 0 | 0 | 0 |
| | RICs | Serum | 0 | 0 | 0 | 0 | 0 | 0 |
| | | BALF | 0 | 0 | 0 | 0 | 0 | 0 |
| | mAbs | Serum | 0 | 0 | 0 | 0 | 0 | 0 |
| | | BALF | 0.29 | 0.29 | 0 | 0.29 | 0.29 | 0 |
| | PBS | Serum | 0 | 0 | 0 | 0 | 0 | 0 |
| | | BALF | 0 | 0 | 0 | 0 | 0 | 0 |
| LIF | RT | Serum | 1.85 | 0 | 2.57 | 0.85 | -0.93 | 1.97 |
| | | BALF | 1.85 | 12.53 | 15.59 | 0.85 | 11.53 | 14.59 |
| | RICs | Serum | 0 | 17.46 | 2.59 | 0 | 0.23 | 2.00 |
| | | BALF | 0 | 16.39 | 1.41 | 0 | 15.39 | 0.41 |
| | mAbs | Serum | 0 | 0 | 3.32 | 0 | -0.93 | 2.83 |
| | | BALF | 0.88 | 0.56 | 0.08 | 0.88 | 0.56 | 0.08 |
| | PBS | Serum | 0 | 14.20 | 0.87 | 0 | 0 | 0 |
| | | BALF | 0 | 0 | 0 | 0 | 0 | 0 |
| IL-13 | RT | Serum | 0 | 0 | 0 | 0 | 0 | 0 |
| | | BALF | 0 | 0 | 0 | 0 | 0 | 0 |
| | RICs | Serum | 0 | 0 | 0 | 0 | 0 | 0 |
| | | BALF | 0 | 0 | 0 | 0 | 0 | 0 |
| | mAbs | Serum | 0 | 0 | 0 | 0 | 0 | 0 |

| | | | | | | | | |
|---|---|---|---|---|---|---|---|---|
| | | BALF | 0 | 10.81 | 0 | 0 | 10.81 | 0 |
| | PBS | Serum | 0 | 0 | 0 | 0 | 0 | 0 |
| | | BALF | 0 | 0 | 0 | 0 | 0 | 0 |
| LIX | RT | Serum | 3534.98 | 4913.72 | 4207.07 | -0.22 | 0.06 | -0.05 |
| | | BALF | 2451.17 | 1883.90 | 1512.10 | 0.51 | 0.31 | -0.05 |
| | RICs | Serum | 4872.74 | 4783.88 | 4396.08 | 0.07 | 0.03 | -0.01 |
| | | BALF | 1778.85 | 2104.48 | 1810.58 | 0.10 | 0.46 | 0.14 |
| | mAbs | Serum | 5154.43 | 4844.09 | 3501.03 | 0.14 | 0.04 | -0.21 |
| | | BALF | 660.02 | 439.03 | 238.66 | -0.59 | -0.69 | -0.85 |
| | PBS | Serum | 4534.86 | 4656.80 | 4443.47 | 0 | 0 | 0 |
| | | BALF | 1618.37 | 1438.32 | 1582.52 | 0 | 0 | 0 |
| IL-15 | RT | Serum | 5.42 | 10.09 | 11.61 | -0.01 | -0.82 | 0.48 |
| | | BALF | 4.51 | 9.72 | 6.03 | 0.31 | -0.65 | -0.13 |
| | RICs | Serum | 9.47 | 20.52 | 11.41 | 0.74 | -0.63 | 0.45 |
| | | BALF | 8.03 | 6.75 | 5.74 | 1.33 | -0.76 | -0.17 |
| | mAbs | Serum | 6.14 | 11.12 | 18.02 | 0.13 | -0.80 | 1.29 |
| | | BALF | 17.56 | 18.32 | 10.66 | 4.09 | -0.35 | 0.55 |
| | PBS | Serum | 5.45 | 55.06 | 7.86 | 0 | 0 | 0 |
| | | BALF | 3.45 | 27.98 | 6.90 | 0 | 0 | 0 |
| IL-17 | RT | Serum | 1.24 | 0.92 | 1.44 | 0.24 | -0.25 | -0.20 |
| | | BALF | 1.30 | 0.88 | 1.10 | 0.30 | -0.28 | 0.10 |
| | RICs | Serum | 1.66 | 0.94 | 1.67 | 0.66 | -0.23 | -0.08 |
| | | BALF | 1.36 | 0.94 | 1.28 | 0.36 | -0.23 | 0.28 |
| | mAbs | Serum | 1.28 | 2.08 | 1.00 | 0.28 | 0.70 | -0.45 |
| | | BALF | 3.97 | 2.44 | 0.89 | 2.97 | 1.00 | -0.11 |
| | PBS | Serum | 1.00 | 1.22 | 1.81 | 0 | 0 | 0 |
| | | BALF | 1.00 | 1.22 | 1.00 | 0 | 0 | 0 |
| IP-10 | RT | Serum | 141.89 | 141.97 | 116.50 | 0.24 | 0.10 | -0.15 |
| | | BALF | 115.87 | 222.41 | 233.21 | 0.49 | 1.59 | 1.59 |
| | RICs | Serum | 239.76 | 172.80 | 190.08 | 1.10 | 0.34 | 0.39 |
| | | BALF | 202.20 | 141.53 | 179.15 | 1.60 | 0.65 | 0.99 |
| | mAbs | Serum | 232.01 | 182.92 | 124.68 | 1.03 | 0.42 | -0.09 |
| | | BALF | 281.46 | 162.95 | 69.79 | 2.63 | 0.90 | -0.23 |
| | PBS | Serum | 114.37 | 128.49 | 137.06 | 0 | 0 | 0 |
| | | BALF | 77.63 | 85.83 | 90.07 | 0 | 0 | 0 |
| | RT | Serum | 135.96 | 437.44*** | 369.34*** | 3.25 | 16.84 | 25.63 |

| | | | | | | | | |
|---|---|---|---|---|---|---|---|---|
| KC (CXCL1) | | BALF | 429.28* | 1830.09*** | 1106.68**** | 4.14 | 36.96 | 39.18 |
| | RICs | Serum | 92.17 | 220.42 | 61.46 | 1.88 | 7.99 | 3.43 |
| | | BALF | 115.66 | 312.52** | 105.68* | 0.39 | 5.48 | 2.84 |
| | mAbs | Serum | 128.18 | 54.38 | 35.87 | 3.00 | 1.22 | 1.59 |
| | | BALF | 366.36 | 303.95 | 136.45 | 3.39 | 5.30 | 3.95 |
| | PBS | Serum | 32.01 | 24.52 | 13.87 | 0 | 0 | 0 |
| | | BALF | 83.46 | 48.21 | 27.54 | 0 | 0 | 0 |
| MCP1 | RT | Serum | 22.29 | 44.80* | 41.33** | 1.21 | 4.65 | 12.53 |
| | | BALF | 34.36* | 68.96** | 60.27*** | 4.14 | 67.96 | 59.27 |
| | RICs | Serum | 39.58 | 15.84 | 26.04* | 2.92 | 1.00 | 7.52 |
| | | BALF | 39.18 | 33.11 | 28.46 | 4.86 | 32.11 | 27.46 |
| | mAbs | Serum | 43.78 | 11.50 | 10.00 | 3.34 | 0.45 | 2.27 |
| | | BALF | 47.21 | 23.28 | 3.54 | 6.06 | 22.28 | 2.54 |
| | PBS | Serum | 10.09 | 7.92 | 3.05 | 0 | 0 | 0 |
| | | BALF | 6.69 | 1.00 | 1.00 | 0 | 0 | 0 |
| MIP-1α | RT | Serum | 0 | 0 | 0 | 0 | 0 | 0 |
| | | BALF | 7.24 | 71.62 | 97.33 | -0.57 | 0.95 | 18.07 |
| | RICs | Serum | 0 | 5.34 | 0 | 0 | 4.34 | 0 |
| | | BALF | 13.11 | 26.05 | 5.10 | -0.22 | 1.30 | 1.48 |
| | mAbs | Serum | 0 | 0 | 0 | 0 | 0 | 0 |
| | | BALF | 284.14 | 155.39 | 80.01 | 15.92 | 12.71 | 37.83 |
| | PBS | Serum | 0 | 0 | 0 | 0 | 0 | 0 |
| | | BALF | 16.80 | 11.33 | 2.06 | 0 | 0 | 0 |
| MIP-1β | RT | Serum | 10.54 | 14.44 | 17.06 | 1.72 | 3.11 | 2.10 |
| | | BALF | 10.54 | 71.62 | 97.33** | 0.32 | 15.31 | 16.67 |
| | RICs | Serum | 24.10 | 11.23 | 6.68 | 5.22 | 2.20 | 0.21 |
| | | BALF | 24.10 | 13.38 | 11.11 | 2.02 | 2.05 | 1.02 |
| | mAbs | Serum | 22.98 | 13.21 | 7.98 | 4.93 | 2.76 | 0.45 |
| | | BALF | 2966.96 | 2071.37 | 124.55 | 370.89 | 470.77 | 21.61 |
| | PBS | Serum | 3.87 | 3.51 | 5.51 | 0 | 0 | 0 |
| | | BALF | 7.98 | 4.39 | 5.51 | 0 | 0 | 0 |
| M-CSF | RT | Serum | 0 | 2.83 | 0 | 0 | 0.18 | -0.17 |
| | | BALF | 0 | 2.83 | 1.31 | 0 | 0.18 | 0.09 |
| | RICs | Serum | 0 | 0 | 0 | 0 | -0.58 | -0.17 |
| | | BALF | 2.83 | 0 | 0 | 1.83 | -0.58 | -0.17 |
| | mAbs | Serum | 1.02 | 1.00 | 1.88 | 0.02 | -0.58 | 0.56 |

| | | | | | | | | |
|---|---|---|---|---|---|---|---|---|
| | | BALF | 3.43 | 5.62 | 5.19 | 2.43 | 1.35 | 3.33 |
| | PBS | Serum | 1.00 | 2.40 | 1.20 | 0 | 0 | 0 |
| | | BALF | 1.00 | 2.40 | 1.20 | 0 | 0 | 0 |
| MIP-2 | RT | Serum | 105.66 | 91.89 | 95.14 | 0.53 | 0.19 | -0.07 |
| | | BALF | 112.25 | 522.59 | 181.66 | 0.90 | 6.19 | 0.95 |
| | RICs | Serum | 103.79 | 98.54 | 109.74 | 0.50 | 0.28 | 0.08 |
| | | BALF | 96.15 | 88.50 | 107.44 | 0.62 | 0.22 | 0.16 |
| | mAbs | Serum | 86.17 | 118.55 | 103.06 | 0.24 | 0.54 | 0.01 |
| | | BALF | 105.85 | 105.42 | 56.88 | 0.79 | 0.45 | -0.39 |
| | PBS | Serum | 69.22 | 77.07 | 101.86 | 0 | 0 | 0 |
| | | BALF | 59.18 | 72.63 | 92.98 | 0 | 0 | 0 |
| MIG | RT | Serum | 837.91 | 702.99 | 585.98 | 0.28 | -0.04 | -0.28 |
| | | BALF | 810.74 | 708.04 | 845.99 | 0.35 | 0.05 | 0.17 |
| | RICs | Serum | 934.12 | 760.25 | 832.78 | 0.43 | 0.04 | 0.03 |
| | | BALF | 873.40 | 720.39 | 775.33 | 0.46 | 0.06 | 0.08 |
| | mAbs | Serum | 798.43 | 819.80 | 1072.62 | 0.22 | 0.13 | 0.32 |
| | | BALF | 175.15 | 392.97 | 303.97 | -0.71 | -0.42 | -0.58 |
| | PBS | Serum | 655.32 | 728.59 | 810.10 | 0 | 0 | 0 |
| | | BALF | 599.96 | 677.42 | 720.85 | 0 | 0 | 0 |
| RANTES | RT | Serum | 4.84 | 6.25 | 4.41 | 0.17 | 0.38 | 0.77 |
| | | BALF | 4.84 | 6.25 | 4.41 | 0.17 | 0.38 | 0.77 |
| | RICs | Serum | 5.70 | 1.47 | 1.53 | 0.38 | -0.67 | -0.38 |
| | | BALF | 5.70 | 1.47 | 1.53 | 0.38 | -0.67 | -0.38 |
| | mAbs | Serum | 4.87 | 3.74 | 4.16 | 0.18 | -0.17 | 0.67 |
| | | BALF | 36.38 | 22.98 | 9.15 | 7.79 | 4.09 | 2.68 |
| | PBS | Serum | 4.14 | 4.52 | 2.48 | 0 | 0 | 0 |
| | | BALF | 4.14 | 4.52 | 2.48 | 0 | 0 | 0 |
| VEGF | RT | Serum | 0 | 0 | 0 | 0 | 0 | 0 |
| | | BALF | 0.73 | 14.87 | 52.57 | -0.60 | 3.07 | 22.43 |
| | RICs | Serum | 0 | 0 | 0 | 0 | 0 | 0 |
| | | BALF | 1.60 | 1.64 | 1.11 | -0.13 | -0.55 | -0.51 |
| | mAbs | Serum | 0 | 0 | 0 | | | |
| | | BALF | 2.23 | 3.30 | 3.46 | 0.21 | -0.10 | 0.54 |
| | PBS | Serum | 0 | 0 | 0 | 0 | 0 | 0 |
| | | BALF | 1.85 | 3.65 | 2.24 | 0 | 0 | 0 |
| TNFα | RT | Serum | 0 | 0 | 0 | 0 | 0 | 0 |

| | | | | | | | |
|---|---|---|---|---|---|---|---|
| | BALF | 0 | 2.81 | 1.31 | -0.59 | 0.23 | 0.31 |
| RICs | Serum | 0 | 0 | 0 | 0 | 0 | 0 |
| | BALF | 0 | 0 | 0 | -0.59 | -0.56 | 0 |
| mAbs | Serum | 0 | 0 | 0 | 0 | 0 | 0 |
| | BALF | 493.91 | 140.86 | 13.27 | 200.18 | 60.54 | 12.27 |
| PBS | Serum | 0 | 0 | 0 | 0 | 0 | 0 |
| | BALF | 2.46 | 2.29 | 1.00 | 0 | 0 | 0 |

[a],MFI determined by Luminex and converted to pg/mL per kit manufacturer instructions; [b], Values are group means (n=6); [c], Fold change relative to mean PBS-treated group concentrations for the respective cytokine, sample source, and time point; *, Significantly different value relative to mean PBS-treated group concentration
